# SKAP binding to microtubules reduces friction at the kinetochore-microtubule interface and increases attachment stability under force

**DOI:** 10.1101/2024.08.08.607154

**Authors:** Miquel Rosas-Salvans, Caleb Rux, Moumita Das, Sophie Dumont

## Abstract

The kinetochore links chromosomes to spindle microtubules to drive chromosome segregation at cell division. We recently uncovered that the kinetochore complex Astrin-SKAP, which binds microtubules, reduces rather than increases friction at the mammalian kinetochore-microtubule interface. How it does so is not known. Astrin-SKAP could affect how other kinetochore complexes bind microtubules, reducing their friction along microtubules, or it could itself bind microtubules with similar affinity but lower friction than other attachment factors. Using SKAP mutants unable to bind microtubules, live imaging and laser ablation, we show that SKAP’s microtubule binding is essential for sister kinetochore coordination, force dissipation at the interface and attachment responsiveness to force changes. Further, we show that SKAP’s microtubule binding is essential to prevent chromosome detachment under both spindle forces and microneedle-generated forces. Together, our findings indicate that SKAP’s microtubule binding reduces kinetochore friction and increases attachment responsiveness and stability under force. We propose that having complexes with both high and low sliding friction on microtubules, making a mechanically heterogeneous interface, is key to maintaining robust attachments under force and thus accurate segregation.

## INTRODUCTION

The kinetochore links chromosomes to spindle microtubules at mitosis. Correct kinetochore-microtubule attachment formation, maturation and maintenance are essential for accurate chromosome segregation. A central function of the kinetochore is to transmit forces generated by motors and microtubules to chromosomes. The kinetochore-microtubule interface must be able to resist these forces to maintain a stable attachment, and yet must be dynamic and responsive to force to track microtubule plus ends and move chromosomes (Rago & Cheeseman, 2013). Work from the last decades has revealed the molecular complexity of the mammalian kinetochore-microtubule interface (Musacchio & Desai, 2017). How the mechanical complexity of this interface arises from its molecular components remains poorly understood. Yet, this is key to understanding chromosome segregation, and its accuracy and failures, as a physical process.

The mammalian kinetochore is composed of about a hundred protein species, each in tens of copies (Johnston *et al*, 2010; Musacchio & Desai, 2017). Three main kinetochore protein complexes play key mechanical roles in microtubule attachments. Ndc80 is a core component of the kinetochore that gets gradually dephosphorylated during mitosis, increasing its affinity for microtubules (DeLuca *et al*, 2005, 2006, 2011; Cheeseman *et al*, 2006; Powers *et al*, 2009; Zaytsev *et al*, 2014, 2015). SKA is instead recruited to bioriented kinetochores, and preserves attachment integrity on depolymerizing microtubules under force (Hanisch *et al*, 2006; Welburn *et al*, 2009; Schmidt *et al*, 2012; Auckland *et al*, 2017; Helgeson *et al*, 2018; Huis In ’T Veld *et al*, 2019). Both Ndc80 and SKA increase friction at the kinetochore-microtubule interface, decreasing dynamics and increasing attachment strength (DeLuca *et al*, 2006, 2011; Zaytsev *et al*, 2015; Long *et al*, 2017; Auckland *et al*, 2017; Helgeson *et al*, 2018; Huis In ’T Veld *et al*, 2019). In contrast, we showed that Astrin-SKAP, recruited to bioriented kinetochores (Fang *et al*, 2009; Schmidt *et al*, 2010; Dunsch *et al*, 2011; Huang *et al*, 2011) appears to decrease friction at the attachment interface, increasing attachment dynamics (Rosas-Salvans *et al*, 2022). SKAP is required for chromosome alignment and segregation (Fang *et al*, 2009; Schmidt *et al*, 2010; Dunsch *et al*, 2011; Wang *et al*, 2012; Huang *et al*, 2011), through its microtubule-binding activity (Friese *et al*, 2016; Kern *et al*, 2017, 2016), but the molecular mechanism by which it decreases friction is not known. For example, SKAP could regulate how other microtubule binders (Ndc80 or SKA) interact with microtubules, reducing their sliding friction on microtubules. Alternatively, and not mutually exclusive, SKAP could bind microtubules with similar affinity but lower sliding friction than other microtubule binders, competing with them for microtubule binding and thereby reducing overall friction. Consistent with these models, SKAP is recruited to kinetochores by Ndc80 (Schmidt *et al*, 2010; Friese *et al*, 2016; Kern *et al*, 2017), and directly interacts with microtubules with a similar binding affinity (Ndc80/Astrin-SKAP) than that of SKA (Ndc80/SKA) (Kern *et al*, 2017; Schmidt *et al*, 2012).

In this work, we ask how SKAP reduces friction at the kinetochore-microtubule interface, combining molecular and mechanical perturbations in human RPE1 cells. We show that SKAP binding to microtubules increases sister kinetochore coordination and decreases tension. Further, kinetochore ablation reveals that SKAP microtubule binding increases attachment sensitivity to forces that trigger kinetochore directional switch. Finally, SKAP binding to microtubules increases attachment stability, preventing detachment under both spindle and microneedle-generated forces. Thus, SKAP’s microtubule binding increases the kinetochore-microtubule interface’s responsiveness and stability under force, essential to accurate segregation. We propose that the presence of low friction components amidst high friction ones, i.e. mechanical heterogeneity, increases the adaptability and thus stability of the kinetochore-microtubule interface. This concept may help us understand other biological interfaces.

## RESULTS

### SKAP_ΔMTBD and SKAP_5D display graded disruption of spindle microtubule binding *in vivo*

To test models for how SKAP reduces friction at the kinetochore-microtubule interface, we sought to perturb SKAP’s spindle microtubule binding *in vivo*. In RPE1 cells, we depleted endogenous SKAP using siRNA (siSKAP) (Dunsch *et al*, 2011; Rosas-Salvans *et al*, 2022), and expressed either wild type (SKAP_WT-GFP) or SKAP mutants with no microtubule-binding activity *in vitro* (SKAP_ΔMTBD-GFP and SKAP_5D-GFP), and with no expected change in Astrin recruitment at kinetochores (Kern *et al*, 2016), essential to recruit SKAP (Schmidt *et al*, 2010; Dunsch *et al*, 2011) (Fig. 1A-C). SKAP_ΔMTBD (88-238aa) is an N-terminus truncation lacking the entire microtubule-binding domain and SKAP_5D contains five point mutations (75-TAT<u>RR</u>NV<u>RK</u>GY<u>K</u>P-87/75-TAT<u>DD</u>NV<u>DD</u>GY<u>D</u>P-87) that decrease the positive charge of the microtubule-binding domain (Fig. 1B) (Kern *et al*, 2016). Live imaging showed, as expected, that SKAP_WT-GFP localizes to bioriented kinetochores, spindle microtubules and spindle poles in metaphase (Fig. 1D) (Kern *et al*, 2016). Both SKAP_ΔMTBD-GFP and 5D-GFP localized to kinetochores, and SKAP_5D’s spindle microtubule and spindle pole localization was stronger than SKAP_ΔMTBD-GFP’s but lower than SKAP_WT-GFP’s (Fig. 1D). This suggests a weak interaction of SKAP_5D with spindle microtubules *in vivo* despite no detectable binding *in vitro* (Kern *et al*, 2016). Thus, SKAP_ΔMTBD and SKAP_5D mutants have impaired yet different microtubule-binding activity in the mitotic spindle in living cells. As such, they are well suited to test the role of SKAP’s microtubule binding in kinetochore-microtubule attachment mechanics.

**Figure 1.**
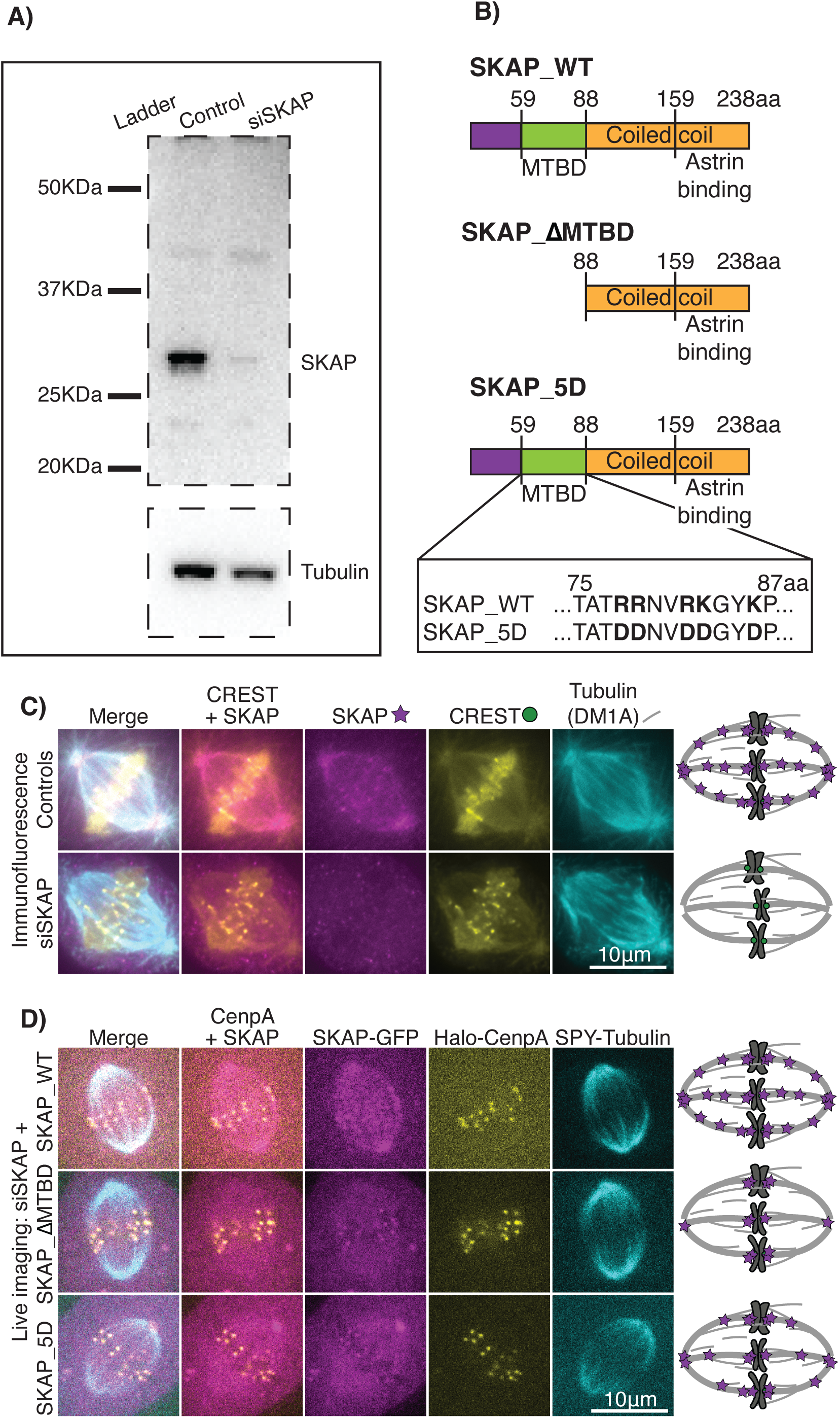
SKAP_ΔMTBD and SKAP_5D display graded disruption of spindle microtubule binding in vivo. **(A)** Representative western blot showing SKAP siRNA efficiency (90% depletion average). **(B)** Schematic representation of SKAP_WT and two mutants expected to disrupt microtubule binding. SKAP_ΔMTBD lacks the first 87 amino acids, including the microtubule-binding domain (MTBD), and SKAP_5D has five point mutations making SKAP’s microtubule-binding domain more negatively charged thus weakening microtubule binding. Coiled coil domains (yellow), including the Astrin-binding domain, microtubule-binding domain (green) and the rest of SKAP’s N-terminus (purple). **(C)** Representative control and siSKAP immunofluorescence images of metaphase RPE1 cells stained for kinetochores (CREST), SKAP and tubulin. Schematic representation of SKAP (magenta) localization at kinetochores (green) and spindle microtubules (grey) (right). **(D)** Representative live images of three above SKAP-GFP constructs in metaphase RPE1 cells under SKAP RNAi. SKAP localizes to kinetochores for all constructs but SKAP_ΔMTBD has much weaker and SKAP_5D weaker spindle microtubule localization than control. Schematic representation of SKAP mutant (magenta) localization at kinetochores and spindle microtubules (grey) (right).

### SKAP binding to microtubules increases sister kinetochore coordination and reduces tension at the kinetochore-microtubule interface

To test the role of SKAP’s microtubule-binding activity at the kinetochore-microtubule interface, we first live-imaged kinetochore oscillations (Fig. 2A) in human RPE1_Halo-CenpA cells (Roscioli *et al*, 2020), labelling kinetochores with Halo tag ligand Janelia Fluor 549 dye and microtubules with SPY_tubulin probe 650 (Dema *et al*, 2022) (Fig. 2B, Supp. Movie 1). Here too, we depleted endogenous SKAP by siRNA and expressed either SKAP_WT-GFP, SKAP_ΔMTBD-GFP or SKAP_5D-GFP. Consistent with our prior work (Rosas-Salvans *et al*, 2022), SKAP depletion decreased kinetochore movement around the plate (control 0.52±0.20μm; siSKAP 0.35±0.18μm; Fig. 2C), speed (control 1.50±0.37μm; siSKAP 1.11±0.21μm; Fig. 2D) and sister velocity correlation (control 0.71±0.12μm; siSKAP 0.60±0.16μm; Fig. 2E) and increased the frequency of opposite sister movement (control 20±6%; siSKAP 26±8%; Fig. 2F) and inter-kinetochore distance (control 1.04±0.08μm; siSKAP 1.30±0.15μm; Fig. 2G), a proxy for tension. SKAP_WT-GFP expression rescued all these SKAP RNAi phenotypes (movement 0.48±0.21μm; speed 1.53±0.37μm; velocity correlation 0.68±0.11μm; opposite movement 22±7%; inter-kinetochore distance 1.08±0.12μm; Fig. 2B-G), consistent with SKAP_WT-GFP being active and the phenotypes specific to SKAP depletion.

**Figure 2.**
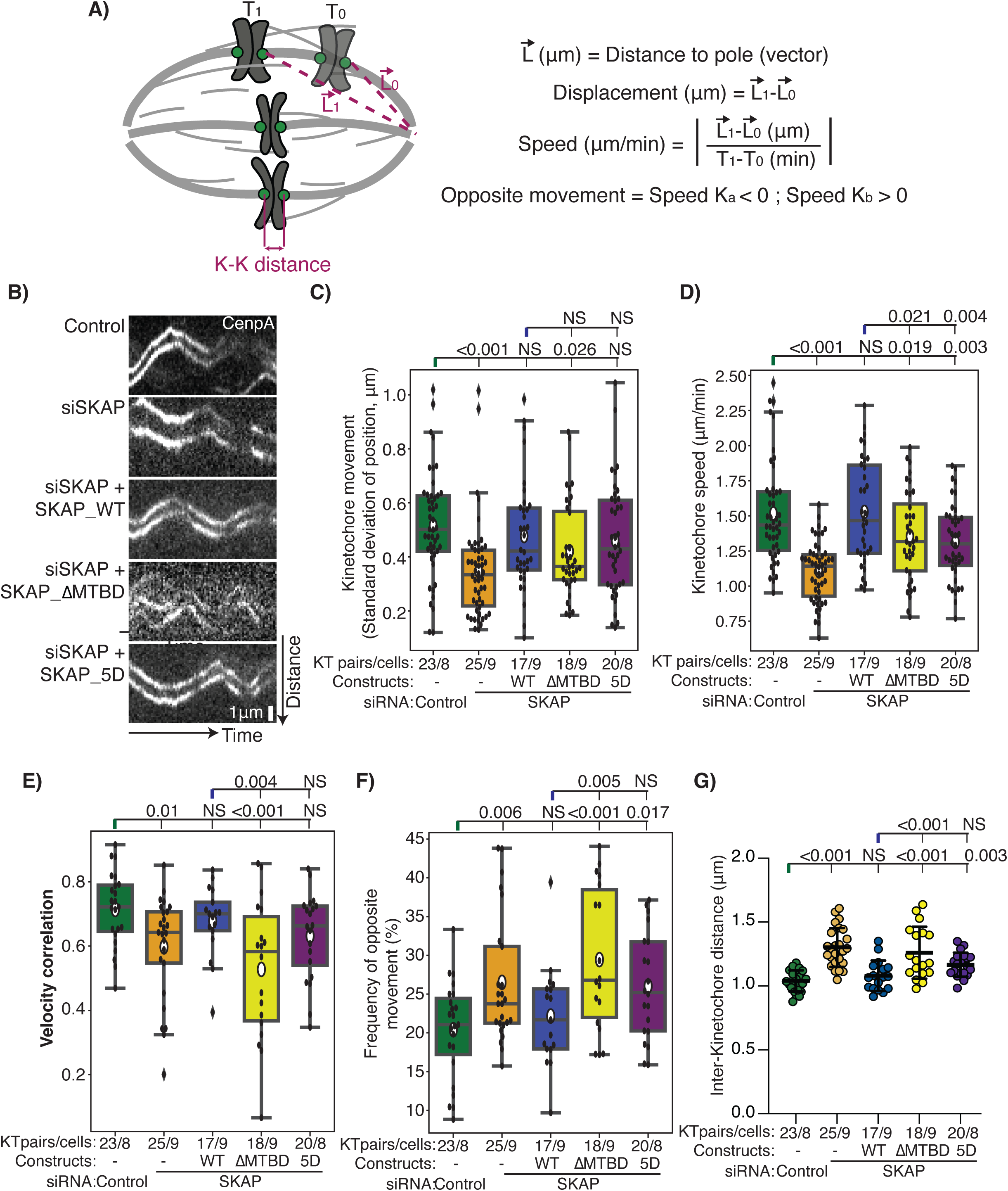
SKAP binding to microtubules increases sister kinetochore coordination and reduces tension at the kinetochore-microtubule interface. **(A)** Cartoon of parameters measured to probe the role of SKAP’s microtubule binding in metaphase kinetochore movement. **(B)** Representative kymographs of sister kinetochore metaphase oscillations timelapse imaging in RPE1 Halo-CenpA (white, JF549) control, siSKAP, and SKAP RNAi cells with either SKAP_WT-GFP, SKAP_ΔMTBD-GFP or SKAP_5D-GFP expression with SPY-tubulin (not shown). Comparison across these five conditions (ANNOVA test) of **(C)** standard deviation of kinetochore position and **(D)** average movement speed of individual kinetochores over time, **(E)** velocity correlation between sister kinetochores, **(F)** fraction of time that individual sister kinetochores move in opposite directions (percentage time) and **(G)** average inter-kinetochore distance for each sister pair. Number of kinetochores (KT) and cells marked for each condition.

In contrast, SKAP_ΔMTBD expressing cells had reduced kinetochore movement (0.42±0.18μm; Fig. 2C), kinetochore speed (1.36±0.32μm; Fig. 2D), and sister velocity correlation (0.53±0.22μm; Fig. 2E) and had increased frequency of opposite sister movement (29±9%; Fig. 2F) compared to SKAP_WT expressing cells or controls. These phenotypes were not dependent on the level of expression of SKAP_ΔMTBD, as they were also present in highly expressing cells (Supp. Fig. 1A-B). SKAP_ΔMTBD expression induced a strong reduction in sister kinetochore coordination, even lower than siSKAP. In contrast, SKAP_ΔMTBD induced only weaker phenotypes associated with kinetochore persistent movement, with movement and speed values significantly lower than controls but also significantly higher than siSKAP. Overall, cells expressing SKAP_5D had less severe phenotypes than SKAP_ΔMTBD cells (movement 0.46±0.21μm; speed 1.32±0.27μm; velocity correlation 0.63±0.14μm; opposite movement 26±7% SD; Fig. 2C-F). Thus, more severe phenotypes were associated with more disruption of microtubule-binding activity, consistent with disrupted microtubule binding being responsible for the observed phenotypes. Further, SKAP_ΔMTBD cells had higher inter-kinetochore tension than SKAP_WT cells (1.26±0.2μm; Fig. 2G), even when highly expressed (Supp. Fig. 1 C). Here too, SKAP_5D cells had an intermediate phenotype (1.17±0.09μm; Fig. 2G), consistent with SKAP’s microtubule binding being essential for kinetochores to respond to the movement of their sister and changes in tension (Fig. 2E-G). The higher inter-kinetochore distance and tension, together with defects in kinetochore movement in SKAP_ΔMTBD cells, suggest that SKAP must interact with microtubules to decrease friction at the attachment interface. Together, the data indicate that SKAP’s microtubule binding increases sister kinetochore coordination and reduces tension at the kinetochore-microtubule interface.

### SKAP’s microtubule binding increases the kinetochore-microtubule interface’s responsiveness to force

Given that SKAP’s microtubule binding increases sister kinetochore coordination and reduces tension at the kinetochore-microtubule interface (Fig. 2), we hypothesized that SKAP binding to microtubules is the key activity required for SKAP’s role in increasing the interface’s responsiveness to force changes (Rosas-Salvans *et al*, 2022). To test this hypothesis, we used a kinetochore ablation assay to probe attachment sensitivity to force changes under SKAP RNAi while expressing either SKAP_WT-GFP or SKAP_ΔMTBD-GFP. In metaphase, kinetochore ablation induces kinetochore-microtubule depolymerization at the remaining sister (Khodjakov & Rieder, 1996; McNeill & Berns, 1981). As that sister approaches the pole, polar ejection force increases (Ke *et al*, 2009), increasing tension at the attachment interface and thereby inducing microtubule rescue and kinetochore directional switch (Fig. 3A) (Khodjakov & Rieder, 1996; McNeill & Berns, 1981; Akiyoshi *et al*, 2010). The kinetochore-microtubule interface’s responsiveness to force determines the distance from the pole at which the kinetochore reverses direction, given an unperturbed polar ejection force gradient (Rosas-Salvans *et al*, 2022).

**Figure 3.**
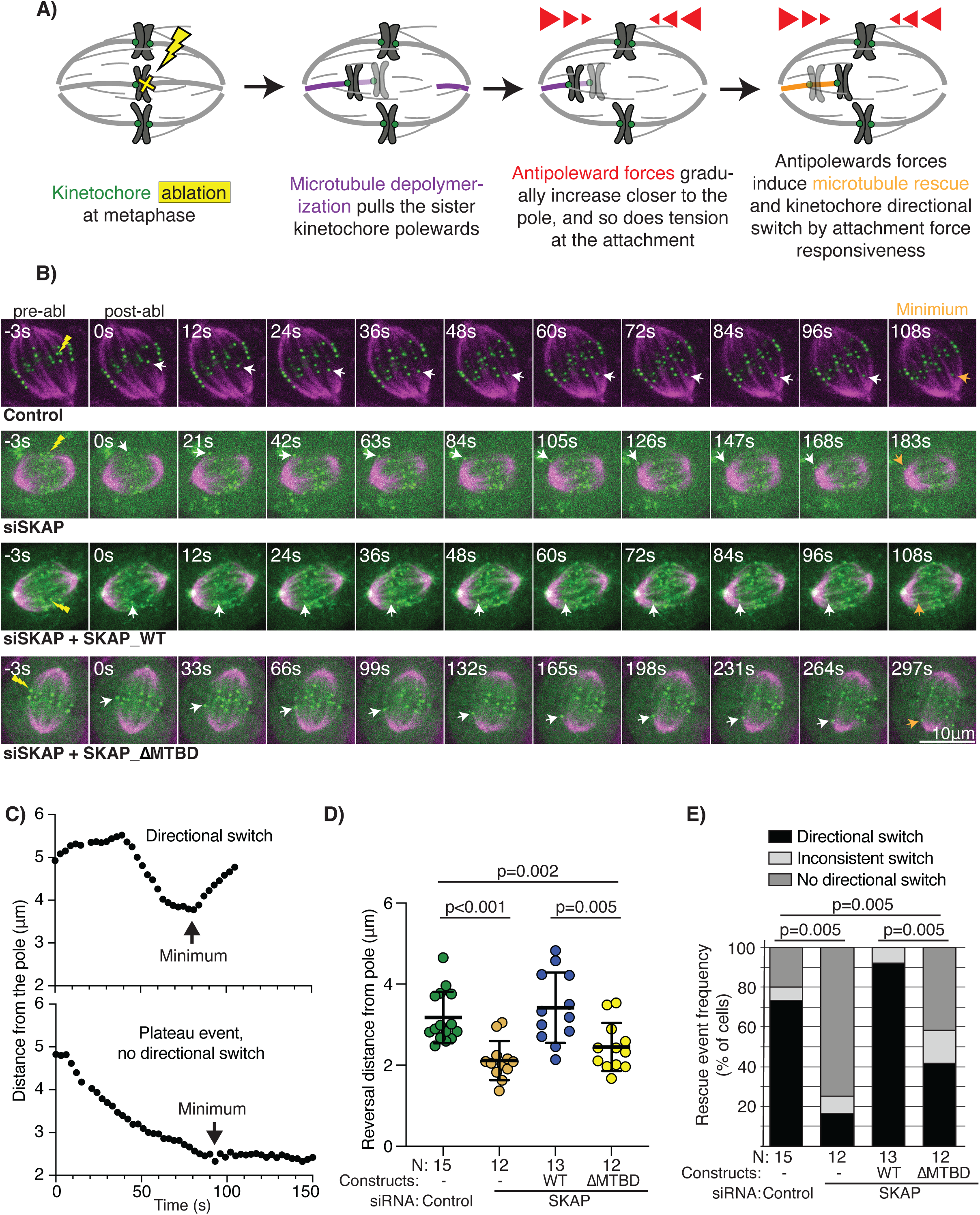
SKAP’s microtubule binding increases the kinetochore-microtubule interface’s responsiveness to force. **(A)** Cartoon of experiment probing the role of SKAP’s microtubule binding in the kinetochore-microtubule interface responding to force: metaphase kinetochore ablation (yellow) reduces tension at the sister kinetochore (green), inducing k-fiber depolymerization (purple) that pulls the sister kinetochore towards the pole, and increasing polar ejection forces (red) approaching the pole induce kinetochore directional switch and k-fiber rescue (orange). **(B)** Representative timelapse images of the above experiment in RPE1 Halo-CenpA (green, Halo tag ligand Oregon Green) cells with SPY-tubulin (magenta) under control, siSKAP, and SKAP RNAi with either SKAP_WT-GFP or SKAP_ΔMTBD-GFP. Timelapse shows kinetochore ablation (yellow bolt, 0 s first frame post-ablation), sister poleward movement (white arrowhead) and subsequent reversal and anti-poleward movement (orange arrowhead) under increasing force. **(C)** Examples of kinetochore movement trajectories defining directional switch and plateau events. For the above cellular conditions, **(D)** kinetochore distance from the pole at the time of directional switch from poleward to antipoleward movement after sister ablation (Mann-Whitney test) and **(E)** percentage of kinetochores with a clear directional switch, no directional switch (plateau) and inconsistent switch near the pole (Fisher exact test). Number of ablation events (N) and cells (also N) marked for each condition.

Ablating single kinetochores, we found that the directional switch of siSKAP kinetochores occurs closer to the pole than for control kinetochores (control 3.18±0.63μm; siSKAP 2.12±0.48μm; Fig. 3B-D, Supp. Movie 2), as expected (Rosas-Salvans *et al*, 2022). This phenotype was rescued by expression of SKAP_WT-GFP (3.42±0.87μm, Supp. Movie 2), confirming specificity, but not rescued by expression of SKAP_ΔMTBD-GFP (2.45±0.59μm) (Fig 3B, D, Supp. Movie 2). Additionally, clear kinetochore directional switches were less frequent in the absence of SKAP or its microtubule-binding activity (Fig. 3C, E, Supp Fig 2A-B). Thus, the microtubule-binding activity of SKAP is key for the kinetochore-microtubule interface’s responsiveness to force changes. Together, the data indicate that SKAP’s binding to microtubules reduces – rather than increases – friction between the kinetochore and microtubules, effectively lubricating the interface and making it more responsive to force.

### SKAP preserves attachment integrity under force through its microtubule-binding activity

Given that SKAP’s microtubule binding increases the kinetochore-microtubule interface’s force responsiveness, we asked whether it helps or hinders the maintenance of robust attachments. While being responsive to force could allow the interface to adapt to force changes by sliding rather than detaching, being too responsive could lead to excess sliding and thereby detachment. Interestingly, inspection of metaphase movies in siSKAP cells expressing SKAP_ΔMTBD (Fig. 2) revealed a putative spontaneous kinetochore detachment event under normal spindle forces. Kinetochore-fiber (k-fiber) microtubules appear to detach from a kinetochore and depolymerize (Supp. Fig. 3 A,B, Supp. Movie 3), as expected without a kinetochore connection (Spurck *et al*, 1990; Maiato *et al*, 2004; Elting *et al*, 2014). Coincidently, the inter-kinetochore distance decreases rapidly (Supp. Fig. 3A,C, Supp. Movie 3), indicating loss of tension and consistent with a detachment event (Auckland *et al*, 2017; Elting *et al*, 2014). We did not observe any such a detachment event in the controls and SKAP_WT expressing cells we imaged. Given that we only observed one putative spontaneous detachment event in SKAP_ΔMTBD cells we could not conclude whether it was due to force on the kinetochore or other factors.

To ask if SKAP’s microtubule binding prevents detachment under force, we sought to externally exert force on a kinetochore-microtubule attachment, and to do so on a kinetochore we select at the time we select. We used microneedle manipulation to apply force in a controlled and reproducible way, and adapted microneedle manipulation of metaphase spindles from rat kangaroo PtK2 cells (Suresh *et al*, 2020; Long *et al*, 2020) to human RPE1 cells. RPE1 cells are rounder and have more chromosomes, making manipulations more challenging. To pull on a k-fiber and its attached kinetochore, we placed the microneedle inside the spindle and against the outermost k-fiber, and moved it perpendicularly to the k-fiber for 10 μm over 18 s. We used RPE1_Halo-CenpA cells, labelling kinetochores with Halo tag ligand Janelia Fluor 549 to monitor the inter-kinetochore distance, and microtubules with SPY_tubulin probe 650 to help direct needle placement. We defined successful manipulations as those that resulted in a large (>0.3 μm) increase of the inter-kinetochore distance, or tension, for the kinetochore pair being pulled on (Fig. 4A-B, Supp. Movies 4-5 and Supp. Fig. 4A-B).

**Figure 4.**
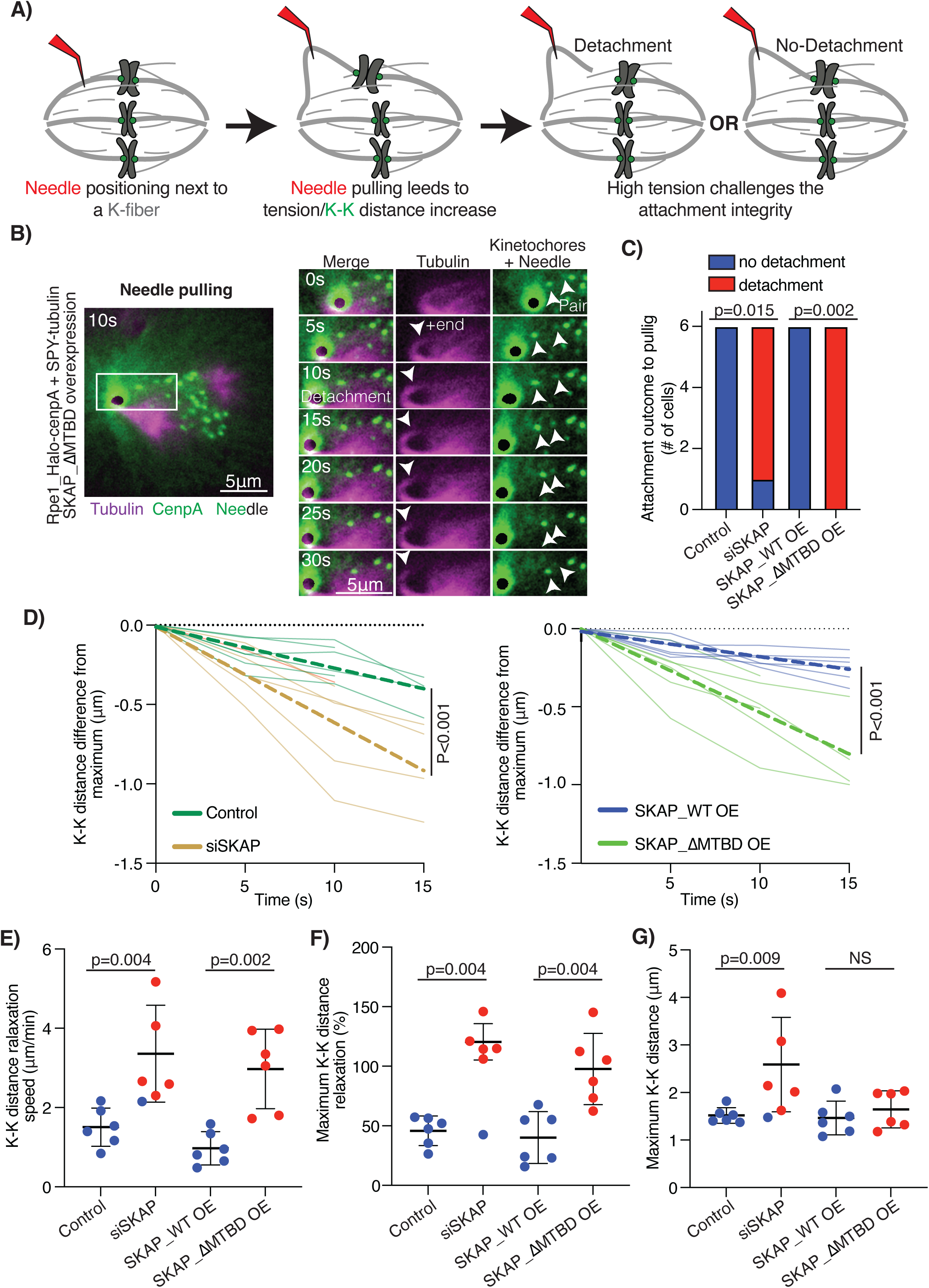
SKAP preserves attachment integrity under force through its microtubule-binding activity. **(A)** Cartoon of experiment probing the role of SKAP’s microtubule binding in attachment robustness under force. A glass microneedle (red) pulls on the k-fiber (grey), increasing tension and inter-kinetochore (green) distance, which leads to either detachment (left) or attachment maintenance (right). **(B)** Representative timelapse images of a kinetochore detachment from its k-fiber in a microneedle (green/black, BSA-Alexa-488) manipulated RPE1 Halo-CenpA control (green, JF549nm, white arrows marking kientochores) cells with SPY-tubulin 650nm (magenta, white arrow marking k-fiber plus-end) expressing SKAP_ΔMTBD-GFP. Whole cell (left) and cropped region (right) timelapse showing increased inter-kinetochore distance (0-10 s) before detachment, the detachment event (between 10 and 15s) and a k-fiber disconnected from the kinetochore and the inter-kinetochore distance decrease after detachment (15-30 s). **(C)** Kinetochore-microtubule attachment outcomes after microneedle pulling in RPE1 Halo-CenpA control, siSKAP, and control cells expressing SKAP_WT-GFP or SKAP_ΔMTBD-GFP (Fisher exact test). Inter-kinetochore distance outcomes for dataset in (C): **(D)** Inter-kinetochore distance relaxation during the first 15 s after maximum inter-kinetochore distance with linear regression fits (dashed lines) for microneedle pulling in control (green) and siSKAP (gold) cells on the left, and control cells expressing SKAP_WT-GFP (blue) and SKAP_ΔMTBD-GFP (green) on the right. **(E)** Average inter-kinetochore distance relaxation over time per kinetochore pair with color indicating detachment (red) vs no detachment (blue) (Mann-Whitney test). **(F)** Inter-kinetochore distance relative relaxation (to final relaxed inter-kinetochore distance) for individual manipulated kinetochore pairs (Mann-Whitney test). **(G)** Maximum inter-kinetochore distance achieved per manipulated pair before relaxation starts (Mann-Whitney test).

Strikingly, upon needle pulling the selected kinetochore detached in all but one siSKAP cells (5/6) and in all the cells overexpressing SKAP_ΔMTBD-GFP (6/6) we manipulated (Fig. 4B-D, Supp. Movie 5). In contrast, we did not observe detachment upon needle pulling in the same number of control cells and in cells overexpressing SKAP_WT-GFP (Fig. 4B-D, Supp. Movie 4). Due to the low throughput nature of these experiments, we opted for overexpression rather than RNAi and rescue to ensure a homogenous molecular background across cells (GFP-labelled SKAP on unperturbed endogenous SKAP level).

To confirm detachments events, kinetochore pairs were qualitatively and quantitatively analyzed upon manipulation. Qualitatively, detachments were determined by detectable loss of connection between the k-fiber and the kinetochore (Supp. Movie 5), concurrent with a sudden decrease of the inter-kinetochore distance, or tension. Quantitative analysis of kinetochore behaviors under needle pulling matched this qualitative analysis. Quantitatively, detached kinetochores showed a more rapid decrease of the inter-kinetochore distance conditions when compared to non-detaching ones, indicating fast tension relaxation by detachment rather than gradual dissipation by microtubule dynamics (Fig. 4D-E). Detached kinetochores also showed higher total and relative relaxation after detachment than non-detached ones (Fig. 4F, Supp. Fig 4C-D). Interestingly, siSKAP kinetochores, but not SKAP_ΔMTBD overexpression kinetochores, reached higher inter-kinetochore distances than controls during pulling (Fig. 4G, Supp. Fig 4B). This suggests that more sliding capacity remains in SKAP_ΔMTBD overexpression than in siSKAP cells, presumably due to different remaining levels of endogenous SKAP. In both cases, fewer SKAP molecules interacting with microtubules decreases attachment stability under force. Thus, SKAP binding to microtubules decreases friction at the attachment interface, increasing both attachment responsiveness and stability under force.

## DISCUSSION

The Astrin-SKAP complex is essential to accurate chromosome segregation (Fang *et al*, 2009; Schmidt *et al*, 2010; Dunsch *et al*, 2011; Huang *et al*, 2011). We recently showed that it decreases friction at the kinetochore-microtubule attachment interface, but how it does – as a microtubule-binding protein – remained unknown. Combining molecular (Fig. 1) and mechanical perturbations in human cells, we show that SKAP’s interaction with microtubules reduces friction at the kinetochore-microtubule interface, increasing its responsiveness to force (Fig. 2-3) and stabilizing it under force (Fig. 4). These findings suggest a model whereby Astrin-SKAP binds microtubules with a similar affinity as other binders, competing with them, but slides with lower friction than them. We propose that a mechanically heterogeneous kinetochore-microtubule interface, with high and low friction elements, adapts more robustly to changes in spindle forces and thus increases attachment force responsiveness and preserves attachment integrity for faithful chromosome segregation (Fig. 5).

**Figure 5.**
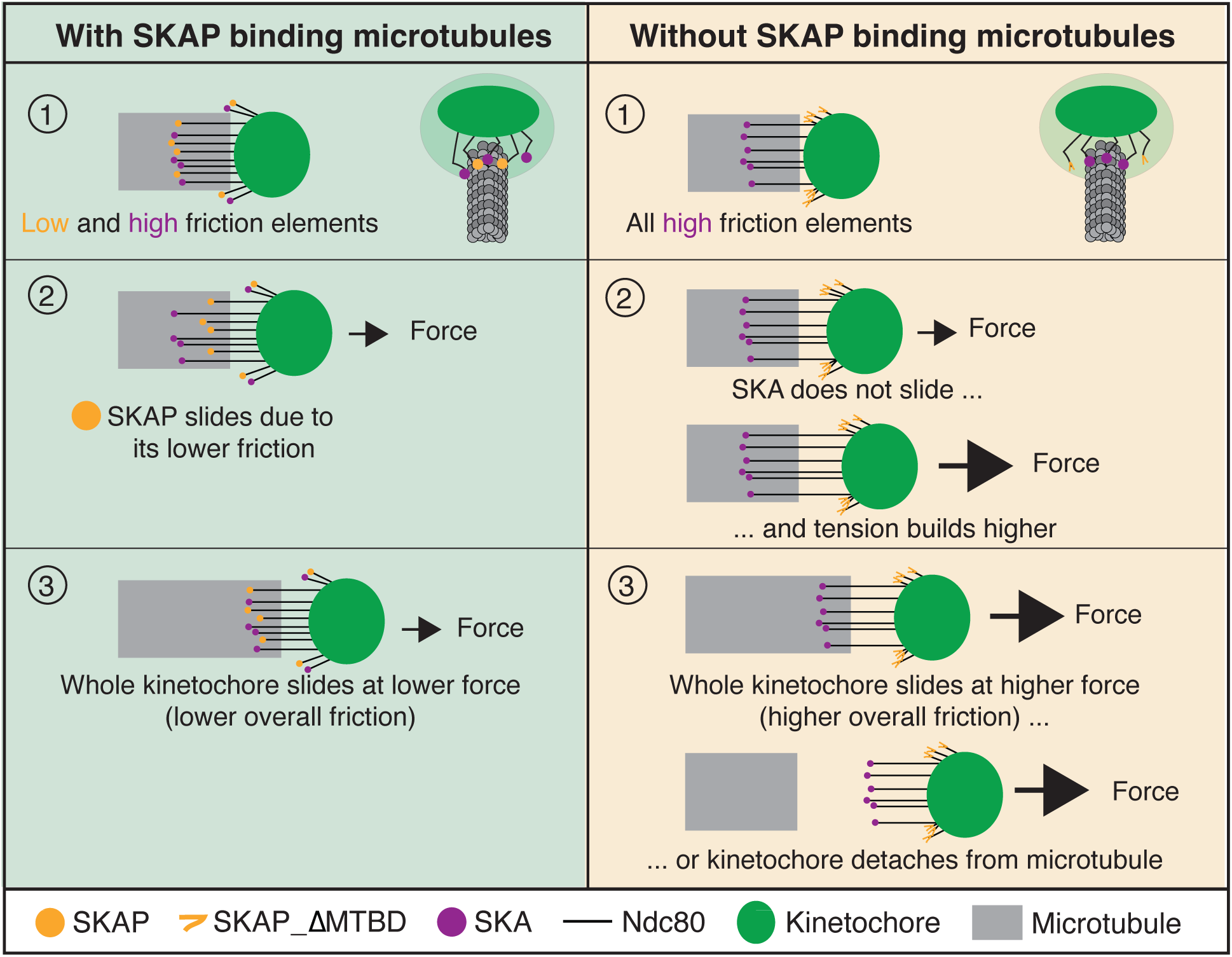
Model for how SKAP’s microtubule binding reduces friction at the kinetochore-microtubule interface, making the attachment more responsive and robust under force. With SKAP (left, 1): 2) Under tension at the kinetochore, SKAP slides before other components due to its lower friction for sliding on microtubules, rearranging the interface and making it responsive to force. 3) This increases tension the other microtubule binders (Ndc80 and SKA) experience, inducing their subsequent sliding. The whole system slides without detachment. Without SKAP or its microtubule-binding activity (right, 1): 2) Under a similar tension as above, there is no rearrangement of the attachment interface since only high friction binders are present. Higher tension is needed to slide components due to their higher friction. 3) Under this higher tension, either the whole system slides and kinetochore and microtubules remain attached or the attachment breaks due to its low rearrangement capacity. SKAP (yellow), SKA (purple), Ndc80 (black), the rest of the kinetochore (green), and microtubules (grey) are depicted.

In principle, SKAP’s microtubule binding could directly regulate the friction needed to slide on microtubules, or regulate microtubule dynamics and indirectly change apparent friction. While future in vitro work will be required to formally uncouple both models, observations we and others made are not easily consistent with the second model: lowering microtubule dynamics with drugs (Waters *et al*, 1998; Okouneva *et al*, 2008) does not recapitulate our observations with siSKAP and SKAP_ΔMTBD (Fig. 2), SKAP depletion does not affect the levels of key microtubule dynamics regulators MCAK and Kif2a (Rosas-Salvans *et al*, 2022), and SKAP’s interaction with EB1 is not necessary for its role in chromosome alignment and segregation (Kern *et al*, 2016; Wang *et al*, 2012; Naoka Tamura *et al*, 2015). Further, changes in attachment mechanics are sufficient to influence microtubule dynamics (Janson *et al*, 2003; Akiyoshi *et al*, 2010).

SKAP’s microtubule-binding activity is essential for kinetochore movement, coordination and tension dissipation (Fig. 2), responsiveness to spindle forces (Fig. 3) and robust attachment under force changes (Fig. 4). Yet, SKAP microtubule-binding mutants don’t have the same magnitude impact on all these mechanical functions. Indeed, both SKAP_ΔMTBD and SKAP_5D have a small impact on parameters associated with kinetochore persistent movement (speed and displacement) (Fig. 2C,D) yet a large impact on kinetochore directional switching (Fig. 3), which affects coordination and tension (Fig. 2E,F). These findings are consistent with a model where a single molecule or mechanical activity cannot account for the full complexity of attachment mechanics. For instance, high frictional components might be more relevant for regulating speed and low frictional ones for directional switch, their combination lead to robust attachments under force. Looking forward, this motivates future work tuning the activities of individual and combinations of attachment factors *in vitro* and *in vivo* to understand how the kinetochore-microtubule interface’s mechanical complexity arises from its molecular parts.

We propose a model in which kinetochore SKAP binds to microtubules with a lower resistance to sliding than other microtubule binders (SKA and Ndc80), yet with similar binding affinity (Kern *et al*, 2017; Schmidt *et al*, 2012). In other words, we propose that SKAP has a lower transition state energy between microtubule lattice sites (ΔG^‡^) but similar free energy of binding (ΔG), possible as both parameters have different structural origins. For example, some proteins even have different frictions (ΔG^‡^) moving towards microtubule minus- vs plus-ends (Forth *et al*, 2014), yet have a single binding affinity (ΔG). In this model, SKAP’s lower friction would help the interface adapt and respond to force, thereby making the attachment stronger under force and avoiding detachment (Fig. 5). With SKAP binding microtubules, an increase in force would induce early sliding of SKAP relative to other binders, locally dissipating force and increasing force on other individual microtubule binders, forcing sliding. This capacity to adapt and react to force changes would result in lower built-up tension (Fig. 2G), higher sister kinetochore coordination (Fig. 2E-F), higher responsiveness to force (Fig. 3) and attachment preservation under force (Fig. 4). In contrast, without SKAP binding microtubules only high friction binders are present at the interface. The same increase in force would not lead to SKAP sliding or higher force on other binders. Thus, higher force on the interface would be needed for its rearrangement. This would lead to higher built-up tension (Fig. 2G), lower sister kinetochore coordination (Fig. 2E-F), lower responsiveness to force (Fig. 3) and detachment under force (Fig. 4) if all or most binders detach at once or produce a zipper-like effect under high tension. Interestingly, *C. elegans* chromosomes lack SKAP, and the gradual recruitment of SKA at their kinetochores freezes chromosome oscillations in metaphase (Cheerambathur *et al*, 2017), consistent with a key role of SKAP maintaining attachment dynamics and force responsiveness. In conclusion, we propose that it is specifically the combination of low and high friction components at the attachment interface that increases attachment adaptability and robustness to force.

How a low friction element makes the kinetochore-microtubule attachment stronger and more robust under force is not known. Notably, the principle of duality of friction components we propose (Fig. 5) has been described in other biological and soft materials, *in silico* and in experiments (Lwin *et al*, 2022; Bayart *et al*, 2016). Adding flexible components to a rigid network significantly increases the strains needed for rupture in cartilaginous tissues (Lwin *et al*, 2022). These tissues have two interpenetrating polymer networks: the stiff network maintains rigidity, while the flexible network distributes mechanical energy via low energy rearrangements, increasing the critical stretch at which rupture begins (Lwin *et al*, 2022). Consistent with this, enhanced sliding in lubricated frictional interfaces increases fracture force (Bayart *et al*, 2016). Both these examples use low-energy movements of soft elements to distribute or dissipate mechanical energy, thereby enhancing resistance to rupture. Biological interfaces often must balance robustness and load-bearing capability with dynamics under force, remaining responsive to force changes. As such, this duality of high and low friction, or rigid and soft components, may help us understand other biological interfaces. Adding lower friction element to an interface may be a general mechanism for increasing the stability and dynamics of this interface, as SKAP does at kinetochore-microtubule attachments.

## ACKNOWLEDGMENTS

We thank Iain Cheeseman for anti-SKAP antibody, Andrew McAnish for RPE1-Halo_CenpA cells, and Andrea Musacchio for discussion. We thank members of the Dumont Lab for discussions and critical reading of the manuscript, particularly Megan Chong and Vanna Tran. This work was supported by a UCSF PBBR Postdoc Independent Research Grant (M.R.S.), NIH R35GM136420 (S.D.) and NSF 1548297 (S.D). S.D. is a Chan Zuckerberg Biohub investigator.

## AUTHOR CONTRIBUTIONS

Conceptualization, M.R.S., M.D., S.D.; Methodology, M.R.S., C.R.; Software, M.R.S.; Validation, M.R.S.; Formal analysis, M.R.S.; Investigation, M.R.S., C.R.; Resources, S.D.; Data curation, M.R.S.; Writing-original draft, M.R.S.; Writing-Review & Editing, M.R.S., C.R., M.D., S.D.; Visualization, M.R.S.; Supervision, M.R.S., S.D.; Funding Acquisition, S.D.

## SUPPLEMENTARY FIGURE LEGENDS

**Supplementary Figure 1 (related to Fig. 2).**
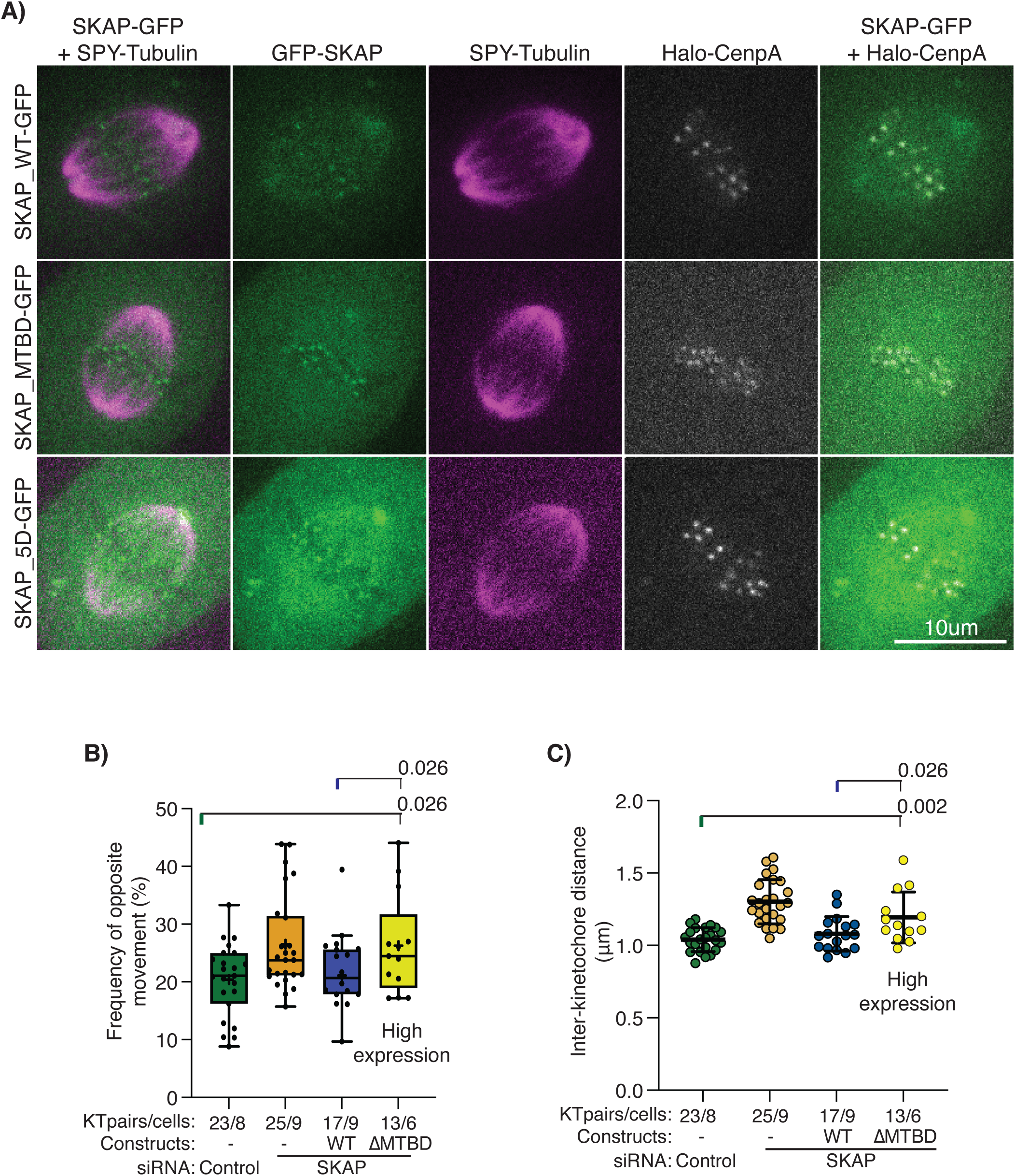
SKAP mutant expression phenotypes are not due to low expression levels. **(A)** SKAP expression levels for cells showed in Fig. 2b. Kinetochore levels of SKAP are similar between conditions. **(B,C)** Cells from Fig, 2 after selecting cells with high expression levels of SKAP_ΔMTBD: **(B)** Fraction of time that individual sister kinetochores move in opposite directions (percentage time) and **(C)** average inter-kinetochore distance for each sister pair between cells.

**Supplementary Figure 2 (related to Fig. 3).**
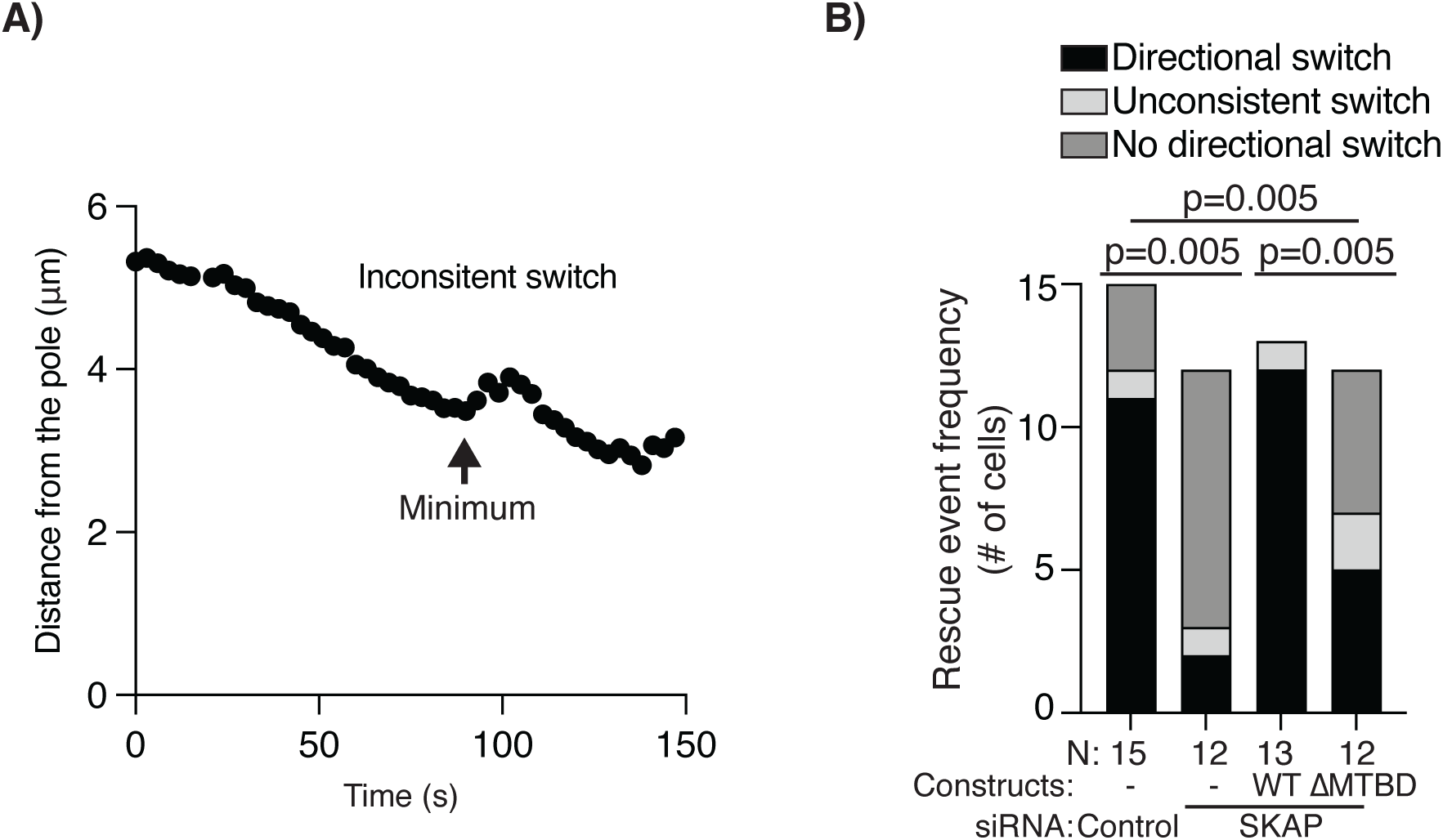
SKAP binding with microtubules increases kinetochore directional switching rate. **(A)** Example of kinetochore movement trajectory defining inconsistent directional switch. **(B)** Percentage of kinetochores with a clear directional switch, no directional switch (plateau) and inconsistent switch near the pole (Fisher exact test). Number of ablation events (N) and cells (also N) marked for each condition.

**Supplementary Figure 3 (related to Fig. 4).**
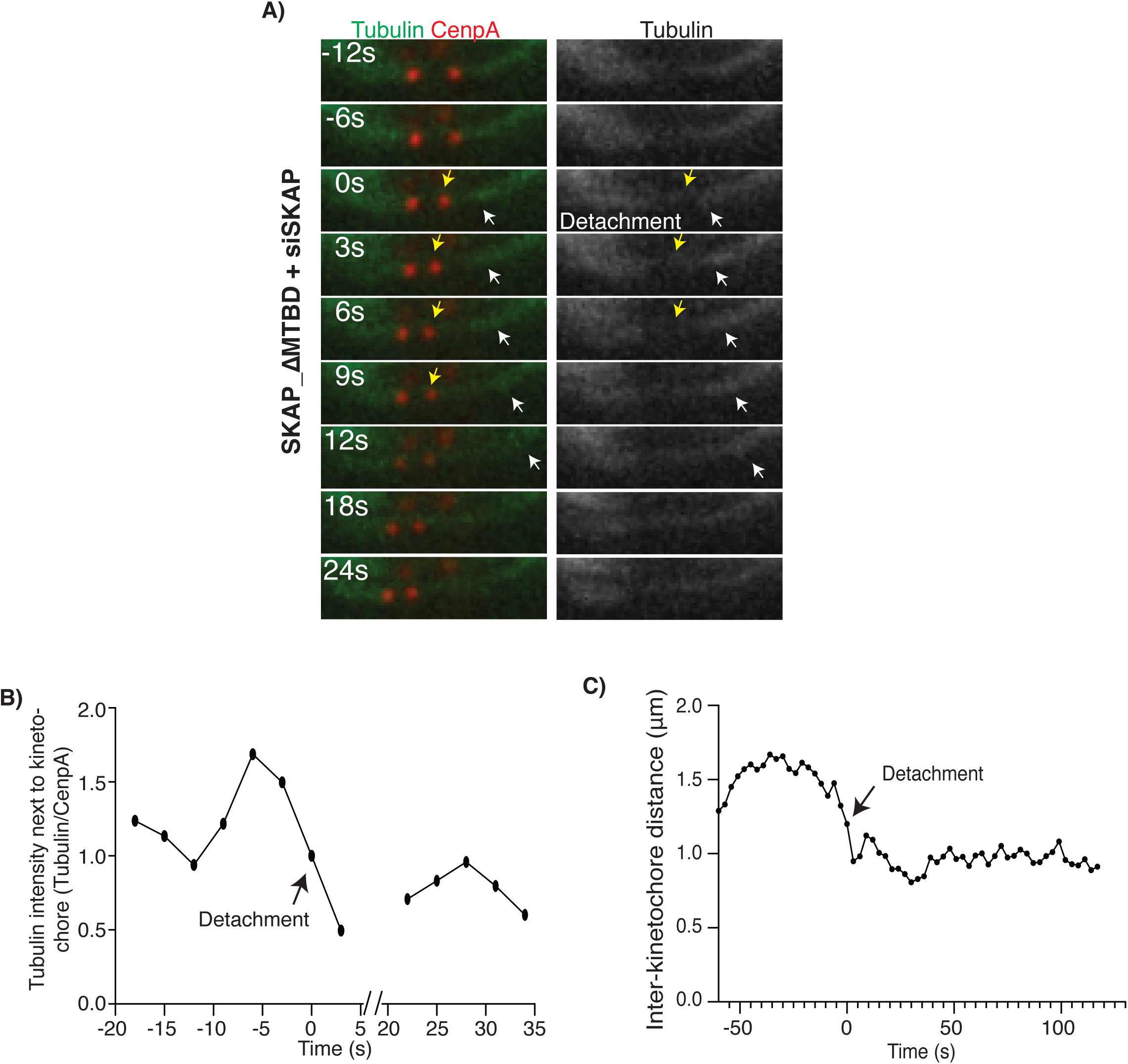
SKAP binding with microtubules increases attachment stability during metaphase oscillations. **(A)** Spontaneous kinetochore detachment during oscillations timelapse. **(B)** Tubulin intensity over time next to the detaching kinetochore in (A). Tubulin intensity normalized to CenpA intensity (to control for plane changes), with timepoints with an unfocused kinetochore not included. **(C)** Inter-kinetochore distance over time of the detaching pair in (A-B).

**Supplementary Figure 4 (related to Fig. 4).**
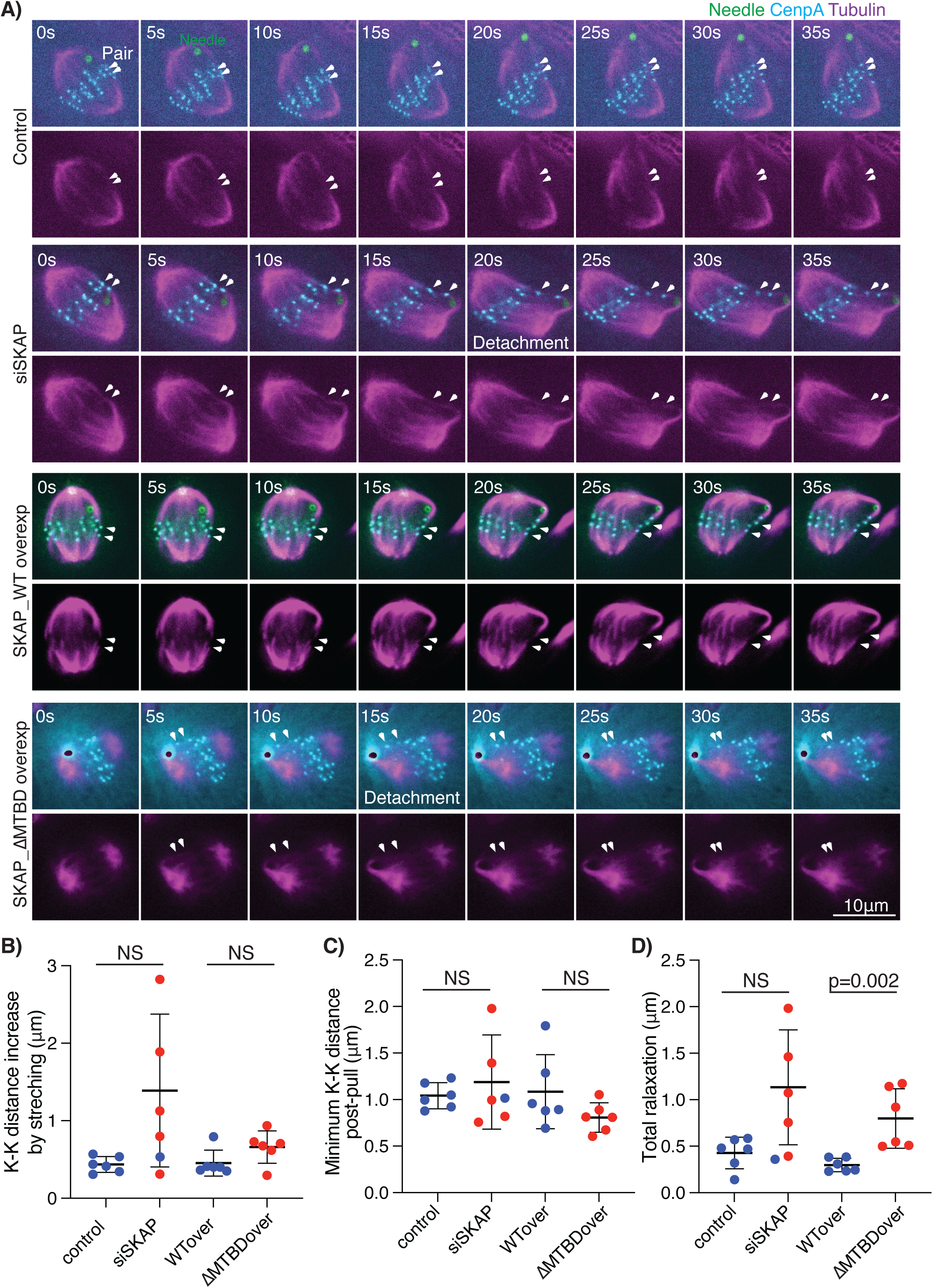
SKAP preserves attachment integrity under force through its microtubule-binding activity. **(A)** Representative microneedle (green or black) manipulation timelapses for RPE1 Halo-CenpA (magenta, JF549nm, white arrows marking kientochores) cells with SPY-tubulin 650nm (magenta) in each condition. For kinetochores that detached (red) and did not detach (blue): **(B)** Inter-kinetochore distance increase due to needle pulling. Mann-Whitney test. **(C)** Minimum inter-kinetochore distance per pair reached after relaxation. Mann-Whitney test. **(D)** Total inter-kinetochore distance relaxation achieved per individual pair. Mann-Whitney test.

## MOVIE LEGENDS

**Supplementary Movie 1 (related to Fig. 2). Kinetochore oscillations in metaphase in cells expressing SKAP_WT-GFP, SKAP_ΔMTBD-GFP or SKAP_5D-GFP in a siRNA background.** Metaphase chromosome oscillations in representative RPE1 Halo-CenpA (green, JF549) with SPY-tubulin (magenta) for each condition: control (top left), siSKAP (top right), siSKAP with SKAP_WT-GFP (bottom left), siSKAP with SKAP_ΔMTBD-GFP (bottom middle), siSKAP with SKAP_5D-GFP (bottom right). Time in min:s.

**Supplementary Movie 2 (related to Fig. 3). SKAP’s microtubule binding increases the kinetochore-microtubule interface’s responsiveness to force.** Representative RPE1 Halo-CenpA (green, Halo tag ligand Oregon Green) cells with SPY-tubulin (magenta) kinetochore ablation timelapses for each condition shown in Fig. 3b: control (top left), siSKAP (top right), siSKAP expressing SKAP_WT-GFP (bottom left) and siSKAP expressing SKAP_ΔMTBD-GFP (bottom right). Time in seconds, with ablation occurring at 0 s. First yellow arrows (t = -3 s) mark the ablated kinetochore, white arrows point to the remaining kinetochore moving polewards, and these become yellow after the directional switch or the minimum value during plateau (siSKAP).

**Supplementary Movie 3 (related to Fig. 4). Attachment disruption in an unmanipulated metaphase cell expressing SKAP_ΔMTBD in a siSKAP background.** Spontaneous detachment event in a RPE1 Halo-CenpA (cyan, JF549) with SPY-tubulin (white) metaphase cell treated with siSKAP and expressing SKAP_ΔMTBD-GFP. Whole cell at the top and kinetochore pair zoom-in at the bottom. White arrow pointing at detached kinetochore after attachment, green arrow pointing to the depolymerizing k-fiber. Time in min:s.

**Supplementary Movie 4 (related to Fig. 4). SKAP preserves attachment integrity under force through its microtubule-binding activity. Micromanipulations of a control cell and a SKAP_WT overexpressing cell.** Representative microneedle manipulation timelapse experiments for a control (left) and a SKAP_WT overexpressing cell (right). Kinetochores in cian, microtubules in magenta and needle in green. Time in min:s. White arrows point to the manipulated kinetochores, moving towards the in the direction of the needle as being pulled and increasing their interkinetochore distance. No detachment detected.

**Supplementary Movie 5 (related to Fig. 4**). SKAP preserves attachment integrity under force through its microtubule-binding activity. Micromanipulation of a siSKAP cell and a SKAP_ΔMTBD overexpressing cell. Representative microneedle manipulation timelapse experiments for RPE1 Halo-CenpA (cyan, JF549nm) cells with SPY-tubulin (magenta) in a siSKAP cell (left) and SKAP_ΔMTBD overexpressing cell (right). The microneedle is in green (siSKAP) or black (SKAP _ΔMTBD). White arrows point to the manipulated kinetochores, moving towards the same direction as the needle as being pulled and increasing their inter-kinetochore distance. Detachment induces fast relaxation of the inter-kinetochore distance as the kinetochore on the needle side moves rapidly away from the needle. Time in min:s.

## METHODS

### Cell culture, siRNA transfection and protein expression in RPE cells

RPE1-Halo_CenpA cells (gift from A. McAinsh, Warwick Medical School) were cultured in DMEM/F12 (Dulbecco’s Modified Eagle Medium: Nutrient Mixture F-12, Thermo Fisher Scientific, 11320082) supplemented with 10% qualified and heat-inactivated fetal bovine serum (FBS, Gibco, 10438-026), penicillin/streptomycin and 300ug/ml Geneticin (G418 sulfate, Gibo, 11811-031), and maintained at 37 °C and 5 % CO_2_. Cells were plated in 35 mm glass-bottom dishes (poly-D-lysine coated; MatTek Corporation) for live imaging experiments or in six-well plates with #1.5 25 mm coverslips (acid cleaned and poly-L-lysine coated) for immunofluorescence or without for immunoblotting. For knockdown and knockdown + expression experiments (Fig. 1-3), siRNA targeting SKAP (5′- AGGCUACAAACCACUGAGUAA-3′) or luciferase siRNA (5’ CGUACGCGGAAUACUUCGA 3’, control) were transfected in above RPE1 cells using 4 µl of lipofectamine siRNAmax (Thermo Fisher Scientific, 13778075) and 10 µM of siRNA in 2 ml of cell culture media. For SKAP rescue and overexpression experiments (Fig. 1-4), 1.5ml of media containing retroviruses were mixed with polybrene (1:1000, EMD Millipore, 4031137) and added to the 2 ml of cell culture media. Cells were incubated at 37 °C for 6-8 h before media wash, and imaged 24 h after transfection and/or infection. SPY-tubulin 555 or 650 (Cytoskeleton Inc., CY-SC203 or CY-SC503) was added to the samples 40 min before imaging at a 1:1000 final concentration from the stock prepared according to manufacturer instructions. Cells were incubated with Halo dyes (Oregon Green (Fig. 3) or Janelia Fluor 549 (Fig. 1,2,4)) for 30 min before imaging.

Retroviral constructs were a gift from I. Cheeseman (Addgene references SKAP_WT 134195; SKAP_MTBD 134194; SKAP_5D 134196 (Kern *et al*, 2016)). For retroviral production Hec293T cells were seeded in a 10 cm dish and transfected with 5 µg of the donor, 4.5 µg of pUVMC and 0.5µg of VSVG using Viafect (Promega, E4982). Cells were incubated for 6 h at 37 °C, followed by a media exchange, and viruses were collected after 24 h incubation with fresh media (twice).

### Immunofluorescence and immunoblotting

SKAP depletion efficiency was validated by immunoblotting and immunofluorescence. For immunoblotting (Fig. 1A), cells were seeded in six-well plates, transfected with Luciferase (control) or SKAP siRNA and processed 24 h after transfection. Cells were collected in PBS1X (Phoshpate Buffered Saline solution) using a cell scrapper and lysed in PBS1X + 1% NP40 on ice for 30 min. Samples were run in a 4-12% Bis-Tris gel (Invitrogen, NPO335BOX) and transferred to a nitrocellulose membrane (Thermo Scientific, 88018). The following primary and secondary antibodies and dyes where used (incubated in TBS1X (Tris-buffered saline), 3 % milk, 0.1 % Tween for 1 h or 45 min, respectively): anti-SKAP (1ug/ml, rabbit, Origene, TA333584), anti-α-tubulin (DM1A, 1:1000, mouse, Sigma, T6199), goat anti-mouse IgG-HRP (1:1000, Santa Cruz Biotechnology, sc-2005,) and mouse anti-rabbit IgG-HRP (1:1000, Santa Cruz Biotechnology, sc-2357). Blots were exposed with SuperSignal West Pico Substrate (Thermo Scientific) and imaged with a Bio-Rad ChemiDoc XRS+ system.

For immunofluorescence experiments cells were fixed in 99.8% methanol for 10 min at -20 °C, 24 h after siRNA transfection and/or infection, and permeabilized in PBS1X, 0.5% BSA, 0.1% Triton (IF buffer thereafter) for 30 min. The following primary antibodies were incubated for 1 h in IF buffer: SKAP (1ug/ml, rabbit, gift from I. Cheeseman (Kern *et al*, 2016)), α-tubulin DM1A (1:1000, mouse, Sigma, T6199), CREST (1:100, human, Antibodies Incorporated, 15-234-0001). Three washes in IF buffer (10 min each) were done before incubation with the following secondary antibodies (1:1000 in IF buffer, 45 min): goat anti-mouse IgG Alexa Fluor 568 (Invitrogen, A11004), goat anti-rabbit IgG Alexa Fluor 488 (Invitrogen, A11008), and goat anti-human IgG Alexa Fluor 645 (Invitrogen, A21445). Samples were washed once in IF buffer and twice in PBS1X (10 min each) before mounting in ProLong Gold Antifade reagent (Thermo Fisher, P36934).

### Microscopy and laser ablation

Samples were imaged using an inverted microscope (Eclipse Ti-E; Nikon) with a spinning disk confocal (CSU-X1; Yokogawa Electric Corporation), head dichroic Semrock Di01-T405/488/568/647 for multicolor imaging, equipped with 405 nm (100 mW), 488 nm (120 mW), 561 nm (150 mW), and 642 nm (100 mW) diode lasers, emission filters ET455/50M, ET525/ 50M, ET630/75M and ET690/50M for multicolor imaging, and an iXon3 camera (Andor Technology) operated by Micro-Manager (2.0.0). Cells were imaged through a 100X 1.45 Ph3 oil objective and 1.5X lens.

For live imaging and laser ablation experiments, cells were maintained in a stage-top incubation chamber (Tokai Hit) at 37 °C and 5 % CO_2_. Metaphase oscillations and ablation experiments were imaged every 3 s (Fig. 1-3). Laser ablation (30-40 pulses of 3 ns at 20 Hz) with 514 nm light was performed using the MicroPoint Laser System (Andor). Successful kinetochore ablation was verified by immediate poleward movement of the remaining sister kinetochore (Fig. 3).

### Microneedle manipulation

Microneedle manipulation was adapted for RPE1 cells based on our previous work in Ptk2 cells (Suresh *et al*, 2020; Long *et al*, 2020). Microneedles were made from glass capillaries with an inner and outer diameter of 1 mm and 0.58 mm respectively (1B100-4 or 1B100F-4, World Precision Instruments). Glass capillaries were pulled with micropipette puller (P-87, Sutter Instruments), bent and polished using a microforge (Narishige International) according to the same specifications, parameters, and geometries described earlier (Suresh *et al*, 2020). These parameters allowed for the needle to approach cells orthogonal to the imaging plane and to conduct manipulations without rupturing the cell (Suresh *et al*, 2020). Prior to imaging, microneedles were coated with BSA-Alexa-488 (Invitrogen, A13100) and BSA-Alexa-555 (Invitrogen, A34786) by soaking them in coating solution for 60 s. Coating solution was obtained by dissolving BSA-Alexa dye and Sodium Azide (Nacalai Tesque) in 0.1 M phosphate-buffered saline (PBS) at a final concentration of 0.02% and 3 mM, respectively (Sasaki *et al*, 2012). This coating solution allows needles to be visualized via fluorescence imaging, aiding in positioning of the needle along a single k-fiber, near to, but not touching, the kinetochore at a z-height appropriate for pulling on k-fibers without overly deforming the membrane. For manipulation experiments, the Tokai Hit incubation chamber lid was removed to allow for needle access. Cells were placed in CO_2_-independent media (Thermo Fisher) and the sample temperature was set at 30 °C (the maximum stable temperature reachable without a lid).

Mitotic cells for microneedle manipulation were chosen based on the following criteria: spindles in metaphase, bipolar shape with both poles in the same focal plane, expressing kinetochore markers, and with high expression of SKAP-GFP constructs when relevant. These criteria were important for pulling on single k-fibers close to the top of the cell with properly bioriented kinetochore pairs that could be tracked for the duration of the manipulation to monitor inter-kinetochore distance and kinetochore-microtubule attachment or detachment.

The micromanipulator was mounted to the microscope body and positioned above samples (Suresh *et al*, 2020). Manipulations were performed in 3D using a x-y-z stepper-motor micromanipulator (MP-225, Sutter Instruments). A 3-axis-knob (ROE-200, Sutter Instruments) was connected to the manipulator via a controller box (MPC-200, Sutter Instruments). Prior to manipulation, the needle was positioned via phase imaging at the approximate x-y position of the intended manipulation site ∼ 10 µm above the cell in z. While imaging every 5 s at three z-planes (0.0 +/- 0.8 um) the needle was manually lowered into place along a single k-fiber, near to, but not touching the kinetochore. If necessary, the needle’s position in x-y was changed by raising the needle above the cell and repositioned in x-y before approaching the intended k-fiber and adjoining kinetochore pair. Once properly positioned, a Python script (Suresh *et al*, 2020) was used to communicate with the Sutter Multi-link software (Multi-Link, Sutter Instruments) to move the microneedle 10 µm over 18 s (62.5 nm resolution) approximately orthogonal to the sister kinetochore-to-kinetochore axis. On occasion, if the spindle body translated greatly during the manipulation and no clear force was transmitted to the sister kinetochores (as seen through a lack of increase in inter-kinetochore distance) a second pull of 5 µm over 9 s was initiated to properly challenge the kinetochore-microtubule attachment. This computer-controlled movement of the microneedle allowed for consistent, reproducible experiments across all studied conditions. Upon completion of needle movement, cells were imaged and the needle remained in place until the stretched kinetochore pair relaxed to a minimum defined by three consecutive timepoints with the kinetochore pairs no longer decreasing in inter-kinetochore distance.

To be analyzed, the manipulated cells had to demonstrate all listed attributes: cell health was not significantly impacted by manipulation (no cell rupture or significant damage to the spindle), the mechanically challenged kinetochore pair was clearly visible throughout the entirety of manipulation and demonstrated clear biorientation, the inter-kinetochore distance of the mechanically challenged pair significantly increased during manipulation (indicating force transmission to the kinetochores), the needle never directly contacted a perturbed kinetochore pair, the cell displayed bright enough tubulin and kinetochore signals to positively identify kinetochore-microtubule detachments. When overexpressing SKAP_WT or mutant SKAP constructs, overexpression had to be easily detectable by the fluorescent signal, and the given construct had to demonstrate proper localization (at kinetochores and spindle poles). To rule out the possibility of k-fiber fracture as opposed to detachment from the kinetochore, tubulin intensity near the suspected detachment site at the k-MT interface was examined.

### Study design and data inclusion criteria

Three general criteria for inclusion of cells in metaphase oscillation, laser ablation and microneedle manipulation experiments were applied. First, cells must express detectable levels of Halo-CenpA at kinetochores, but not so high as to completely label chromosome arms. Second, cells must be in metaphase, with the chromosomes aligned at the center of the spindle forming a defined metaphase plate. Third, SKAP expressing cells must show clear detectable SKAP-GFP at kinetochores, with comparable intensity between conditions. Due to the substantially different localization of the mutants, overall expression levels per cell were not measured. Instead, SKAP-GFP levels at kinetochores were visually used as reference for expression level. An extra analysis including exclusively highly expressing cells was done to ensure the observed changes were due to the specificity of the mutant phenotypes instead of differences in expression levels between mutants (Supp. Fig. 1). For oscillation experiments (Fig. 1-2), 2-4 kinetochore pairs per cell were analyzed, and both kinetochores from the pair must stay in focus for a minimum of 180 s. For kinetochore ablation experiments (Fig. 3), ablations where the sister kinetochore did not react, i.e. did not consistently move polewards, were excluded. For microneedle manipulation experiments (Fig. 4), cells were excluded in the absence of detectable increase of the inter-kinetochore distance upon needle pulling and if the pulled sister pair went out of focus right before or during pulling, or before full relaxation.

We did not pre-estimate a required sample size before performing experiments nor did we blind or randomize samples during experimentation or analysis. The ablation and microneedle manipulation experiments in this study are low throughput by nature, which does not enable us to report averages from multiple independent replicate experiments. Instead, we pool cells from across different independent experiments (with at least three independent experiments per condition per assay).

### Analyzing kinetochore behaviors

For metaphase oscillations and ablation experiments, cells where processed and analyzed in FIJI. Timelapse images were registered using the MultiStackReg plugin and kinetochores and poles were manually tracked from Halo-CenpA and SPY-tubulin movies using the MtrackJ plugin. The center of the spindle pole was used as a fixed position for the pole reference. Kinetochore position was calculated as the distance from the spindle pole position. In metaphase oscillations experiments, all quantifications were performed using home-written Python code (deposited in github.com). Kinetochore movement was calculated by obtaining the standard deviation of the position of each individual kinetochore. Kinetochore speed at each timepoint was calculated as the difference in kinetochore position between two consecutive timepoints (Fig. 2). Sister kinetochore movement coordination was obtained by calculating the correlation of sister kinetochore velocity over time (Fig. 2E) or by the percentage of timepoints in which sister kinetochore movement direction was opposite (Fig. 2F). Inter-kinetochore distance was calculated by obtaining the vector linking the centroids of the two sister kinetochores, obtaining the length of the vector.

In ablation experiments, time after ablation was measured from the fist timepoint immediately after ablation. A kinetochore directional switch was defined by the first relative minimum at the kinetochore track graph (Fig. 3C,D). Rescue event frequency was analyzed by quantifying the presence or absence of a visually clear and consistent kinetochore directional switch on the kinetochore tracking graphs (Fig. 3C,E, Supp. Fig. 2A,B).

In microneedle manipulation experiments, inter-kinetochore distance was obtained by calculating the distance between the two centroids of the kinetochore pair. Inter-kinetochore distance relaxation was obtained by calculating the slope of the linear regression line for each individual pair (Fig. 4D,E). Total inter-kinetochore distance relaxation was obtained by subtracting the minimum inter-kinetochore distance post-stretch from the maximum inter-kinetochore distance during the pull (Supp. Fig. 4D), while the percentage was calculated relative to the minimum (Fig. 4F). Inter-kinetochore distance increase by stretching was obtained by subtracting the minimum inter-kinetochore distance pre-pull from the maximum inter-kinetochore distance during pulling (Supp. Fig. 4B).

### Movie preparation

Movies were prepared using FIJI. Brightness and contrast were linearly adjusted to clearly visualize the kinetochores and centrioles.

### Statistical analysis

Statistical analysis was performed in Python or Graphpad (Prism 9). The Fisher’s exact test was used in for qualitative data comparisons (Fig. 3E and 4C). ANOVA test was used for multiple comparisons (Fig. 2C-G, Supp. Fig. 1B,C). Mann-Withney test was use for paired comparisons of continuous variables (Fig. 3D, 4E-G, Supp. Fig. 4B-D). Analysis of covariance test, ANCOVA, was used for linear regression slopes comparison (Figure 4D,E). Unless otherwise noted, results in the text represent mean ± standard deviation.

